# RAF conformational autoinhibition and 14-3-3 proteins promote paradoxical activation

**DOI:** 10.1101/849489

**Authors:** Gaurav Mendiratta, Kodye Abbott, Yao-Cheng Li, Jingting Yu, Jianfeng Huang, Maxim N. Shokhirev, Thomas McFall, Geoffrey M. Wahl, Edward C. Stites

## Abstract

RAF kinase inhibitors can, in some conditions, increase RAF kinase signaling. This process, which is commonly referred to as “paradoxical activation” (PA), is incompletely understood. RAF kinases are regulated by autoinhibitory conformational changes, and the role of these conformational changes in PA is unclear. Our mathematical investigations reveal that a dynamic equilibrium between autoinhibited and non-autoinhibited forms of RAF, along with the RAF inhibitor stabilization of the non-autoinhibited form, can be sufficient to create PA. Using both computational and experimental methods we demonstrate that 14-3-3 proteins, which stabilize both RAF autoinhibition and RAF dimerization, potentiate PA. Our model led us to hypothesize that increased 14-3-3 expression would amplify PA for the third generation RAF inhibitors that normally display minimal to no PA. Our subsequent experiments find that 14-3-3 overexpression increases PA, increases RAF dimerization, and promotes resistance to these inhibitors, effectively “breaking” these “paradox breaker” and pan-RAF inhibitors. Overall, this work reveals a robust mechanism for PA based solely on equilibrium dynamics of canonical interactions in RAF signaling and identifies conditions which allow PA to occur.

## Introduction

The RAS/RAF pathway plays essential roles in human cancer. Proliferation signals generated by transmembrane receptors signal through RAS GTPases to the RAF kinases that initiate the RAF/MEK/ERK Mitogen Activated Protein Kinase (MAPK) cascade. Mutations within this pathway are very common in cancer ^1^. Multiple drugs have been developed to inhibit the RAF kinases ^2, 3, 4, 5^ and these agents have proven clinically valuable for melanoma ^6, 7, 8^ and colorectal cancer ^9, 10, 11^. Additionally, RAF inhibitors appear promising for other BRAF mutant cancers ^12, 13, 14^.

As the RAF kinases (BRAF, CRAF, and ARAF) are key conduits of signals from the RAS GTPases (KRAS, NRAS, and HRAS), it was originally hoped that RAF inhibitors would be able to block the transmission of RAS signals (**Fig. 1a**). However, RAF inhibitors were instead unexpectedly found to amplify RAS signals through a process that is commonly referred to as “paradoxical activation” (PA) (**Fig. 1b**) ^15, 16, 17^. Despite numerous studies, the mechanisms driving PA are still not fully understood ^4, 18, 19^. The regulation of RAF kinase activation is complex with multiple regulatory steps ^18, 20, 21, 22^, and several of these processes have been claimed to play a role in PA ^15, 17, 18, 23, 24, 25, 26^.

**Fig. 1:**
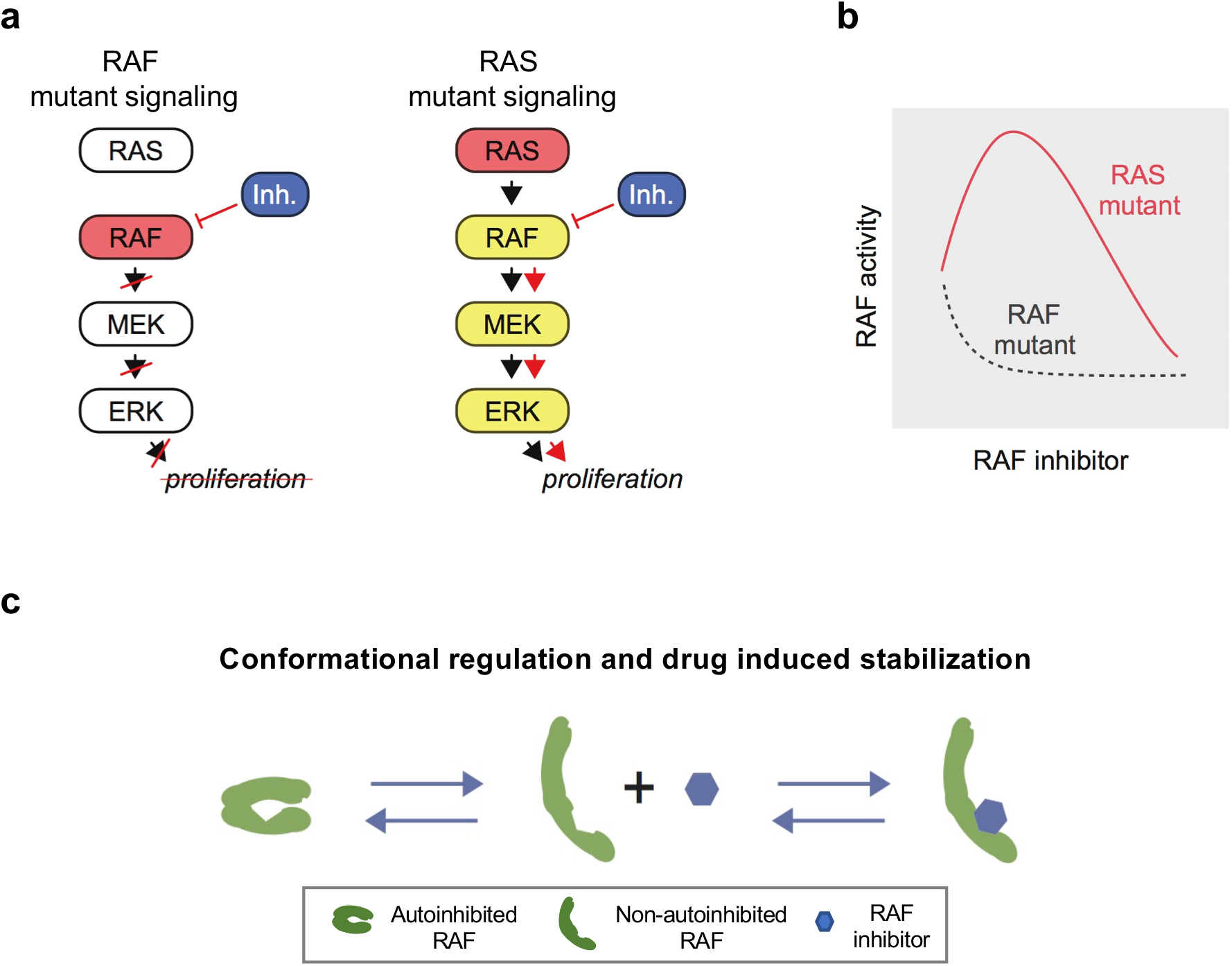
Paradoxical activation (PA) and possible mechanisms. **a** Schematic of the RAS/RAF/MEK/ERK signaling pathway. Intuition for this pathway originally posited that a RAF inhibitor would be able to inhibit signaling for both RAF and RAS mutant cancers. **b** Schematic of the PA concept. When a RAF inhibitor is given to a RAF mutant cell, there is a progressive decline in RAF signaling. However, when given to a RAS mutant cell, there is an increase in RAF signaling at lower doses of RAF inhibitor and suppression of RAF signaling at higher doses of inhibitor. **c** RAF autoinhibition and the stabilization of the non-autoinhibited, dimerization and signaling competent form of RAF by inhibitor has been proposed to contribute to PA, and is here analyzed mathematically to investigate possible implications.

Conformational changes of the RAF monomer contribute significantly to RAF kinase activation (**Fig. 1c**) ^20, 22, 27, 28^. In the “autoinhibited” form, associations between its N-terminus and its kinase domain maintain RAF in an inactive form that does not dimerize ^20, 29^. In the “non-autoinhibited” form, the kinase domain is no longer occluded, and other regulatory mechanisms that contribute to full RAF kinase activation may occur, such as kinase domain conformational changes and dimerization ^20, 30^. Recent experimental work reports that RAF inhibitors tend to promote a net transition to the non-autoinhibited conformation that is bound to RAS-GTP ^19, 27^. It has previously been suggested that this biasing to the non-autoinhibited state may contribute to PA ^27^.

Here, we report our mathematical analysis of how RAF autoinhibitory conformation changes drive PA and the experimental testing of a surprising insight from the modeling that has important clinical implications. On the theoretical side, we developed a series of mathematical models that describe key regulatory steps that have been implicated in RAF signaling. We developed these models to follow biochemical and thermodynamic principles, and we derive the behaviors that logically follow from these principles and mechanisms. Our modeling reveals that RAF autoinhibitory conformational changes alone can be sufficient to drive PA, and we derive the conditions where RAF autoinhibitory conformation drives PA (and also where this same system will fail to induce PA). We extended our model to include the roles of 14-3-3 proteins in stabilizing RAF in the autoinhibitory state and also in stabilizing RAF dimers ^25, 28, 31^, ^32, 33^. We find 14-3-3 can further potentiate PA. This led us to hypothesize that drugs developed to display minimal to no PA under standard cellular conditions could be compromised by increased PA due to amplified 14-3-3 expression. Our experiments confirm this, revealing that 14-3-3 overexpression can amplify PA for existing RAF inhibitors and can even create PA for third generation RAF inhibitors that were developed not to display PA. Our experiments characterize the increase in dimerization from 14-3-3 expression that accompanies PA, and also suggest increased 14-3-3 expression may be a mechanism of resistance in terms of increased off-target effects should these new RAF inhibitors make it to the clinic.

## Results

### Analytical modeling of RAF autoinhibition

A mathematical model of biochemical processes allows one to rigorously analyze what behaviors are possible for a given set of reaction mechanisms and can lead to non-obvious hypotheses for further experimental testing ^34, 35, 36, 37, 38, 39^. We developed a mathematical model to study whether the stabilization of RAF in its non-autoinhibited state by RAF inhibitors may be sufficient to generate PA. Our first mathematical model allows RAF to adopt two different conformations: one of which is autoinhibited and can neither dimerize nor bind drug, the other of which is non-autoinhibited and can bind drug and/or dimerize ^30^. Drug-bound RAF is assumed to only be able to transition back to an autoinhibited state after any bound drug has dissociated. Within the model, wild-type RAF is implicitly assumed to be activated by RAS-GTP, as binding to RAS-GTP is an essential step to wild-type RAF activation ^20^. A complete mechanism is portrayed in **Fig. 2a**. We derived the steady-state solution for this system using the principal of detailed balance ^37, 38^. We focus on steady-state solutions because PA is a steady-state phenomenon that is reflected in the long-term outgrowth of secondary tumors with RAS mutations from patients treated with a RAF inhibitor^40^.

**Fig. 2:**
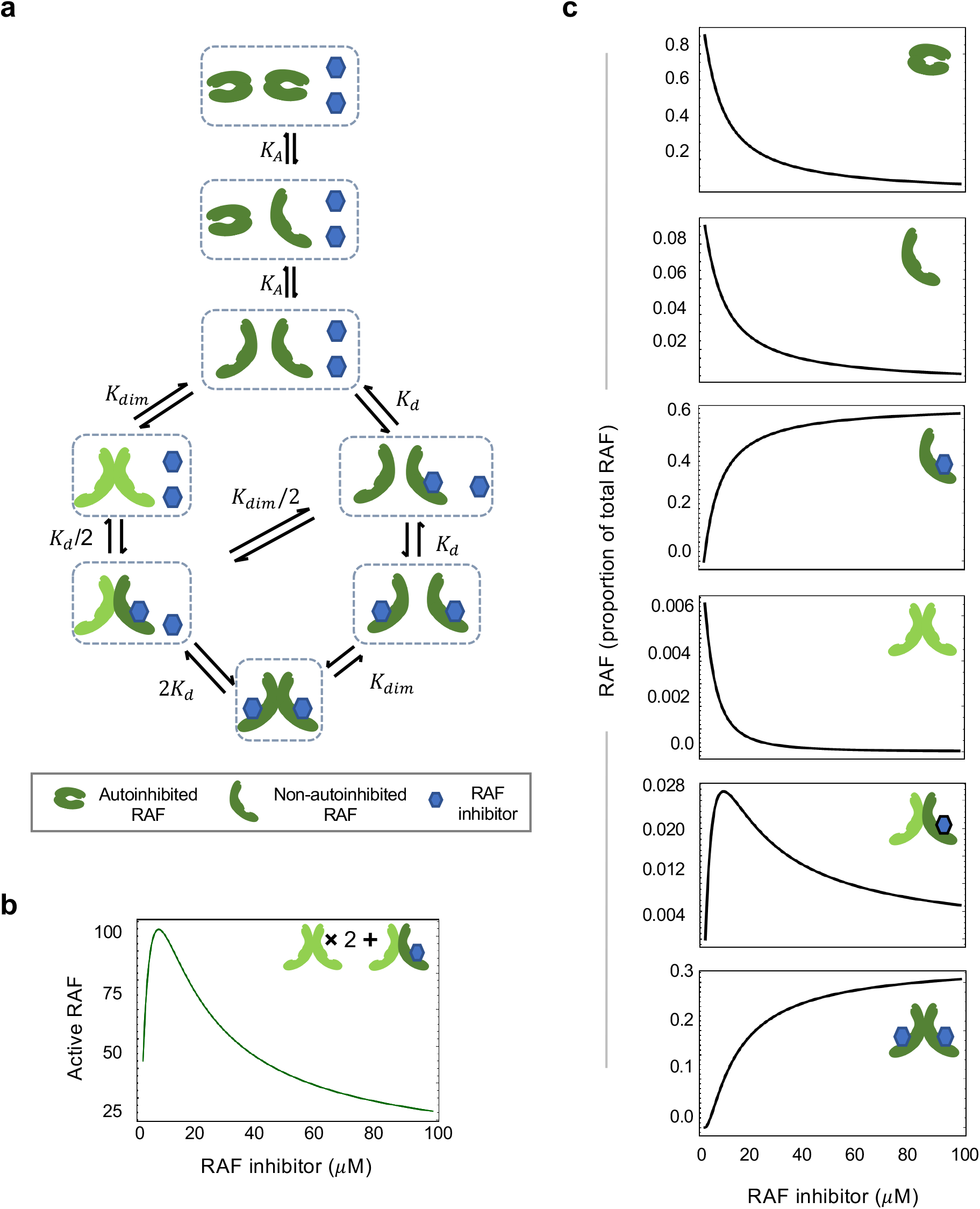
RAF autoinhibition is a mechanism that can produce paradoxical activation. **a** Schematic of the RAF autoinhibition and dimerization model. **b** Representative plot demonstrating that this mechanism is sufficient to generate paradoxical activation. Plotted quantity is the number of active RAF protomers (within a dimer, not bound to drug) as a function of RAF inhibitor abundance, normalized to the maximum. **c** Representative plots portraying the proportion of total RAF in its different states to illuminate the signaling state changes underlying PA.

### Paradoxical activation is a robust outcome of conformational autoinhibition

We investigated whether, and when, the mechanism in our model was sufficient to create PA. We can demonstrate that the presence of both conformational autoinhibition and stabilization of the active form by RAF inhibitors is sufficient to create PA for a wide range of system parameters. Additionally, our analysis yields the conditions necessary for PA to occur by this mechanism. In other words, PA does not always happen when conformational autoinhibition is present. We can predict PA for a wide range of RAF concentration values and visualize the range of system parameters for which conformational autoinhibition promotes PA (**Fig. 2b, Supplementary Fig. 1**).

### Paradoxical activation reflects a shifting balance of signaling complexes

To understand and illustrate how PA arises from this mechanism, we considered the proportion of RAF in each of its possible states: (i) autoinhibited RAF monomer, (ii) non-autoinhibited RAF monomer that is not bound to drug, (iii) non-autoinhibited monomer that is bound to drug, (iv) RAF dimer with no drug bound, (v) RAF dimer with one of two kinase domains bound to drug, (vi) RAF dimer with both kinase domains bound to drug. We considered the total amount of kinase activity to be the number of RAF protomers within a dimer that are not bound to drug. An essential role of dimerization for wild-type RAF activation is supported by prior work ^20, 21^. As the RAF inhibitors described are ATP competitive ^41, 42^, there can be no kinase activity within the dimer when both protomers are bound to inhibitor. However, a RAF dimer is believed to be capable of signaling when only one of the two protomers is bound to drug ^18, 19^.

Before drug is given, a significant fraction of RAF is autoinhibited and there are low levels of non-autoinhibited RAF and RAF dimers. This initial condition is chosen as our analysis shows this is necessary for PA to be generated via conformal autoinhibition based mechanism. As RAF inhibitor levels increase, the level of autoinhibited RAF progressively declines (**Fig. 2c**). Non-autoinhibited RAF distributes between drug-bound monomeric and dimeric forms while the unbound monomeric form maintains equilibrium with the autoinhibited RAF (**Fig. 2c**). The increased quantity of RAF dimers reflects the increased pool of RAF proteins that are non-autoinhibited and therefore capable of dimerization. This results in a drug-dependent increase in RAF dimers bound to drug in one site (**Fig. 2c**) and thereby the increase in total RAF kinase activation that accounts for PA. The quantity of drug-bound RAF monomer and doubly-drug-bound RAF dimer progressively increases to saturation as a function of the drug amount, resulting in the eventual reduction in RAF kinase activity that is associated with PA dose responses (**Fig. 2c**).

### 14-3-3 proteins are mathematically predicted to be capable of increasing the range, magnitude, and potential for paradoxical activation

We next asked how proteins that stabilize the autoinhibited form of RAF might influence PA. 14-3-3 proteins are phosphoserine binding proteins that interact with a large number of proteins and contribute to a wide variety of cellular processes ^43^. The RAF kinases are well-known binding partners for 14-3-3 ^44^. 14-3-3 proteins bind and stabilize the autoinhibited form of BRAF ^28, 31^, ^32, 33, 45^. However, 14-3-3 also stabilizes the RAF dimers ^25, 31^, ^32, 42, 46, 47^. We extended our model to include both of these sets of reactions (**Fig. 3a**).

**Fig. 3:**
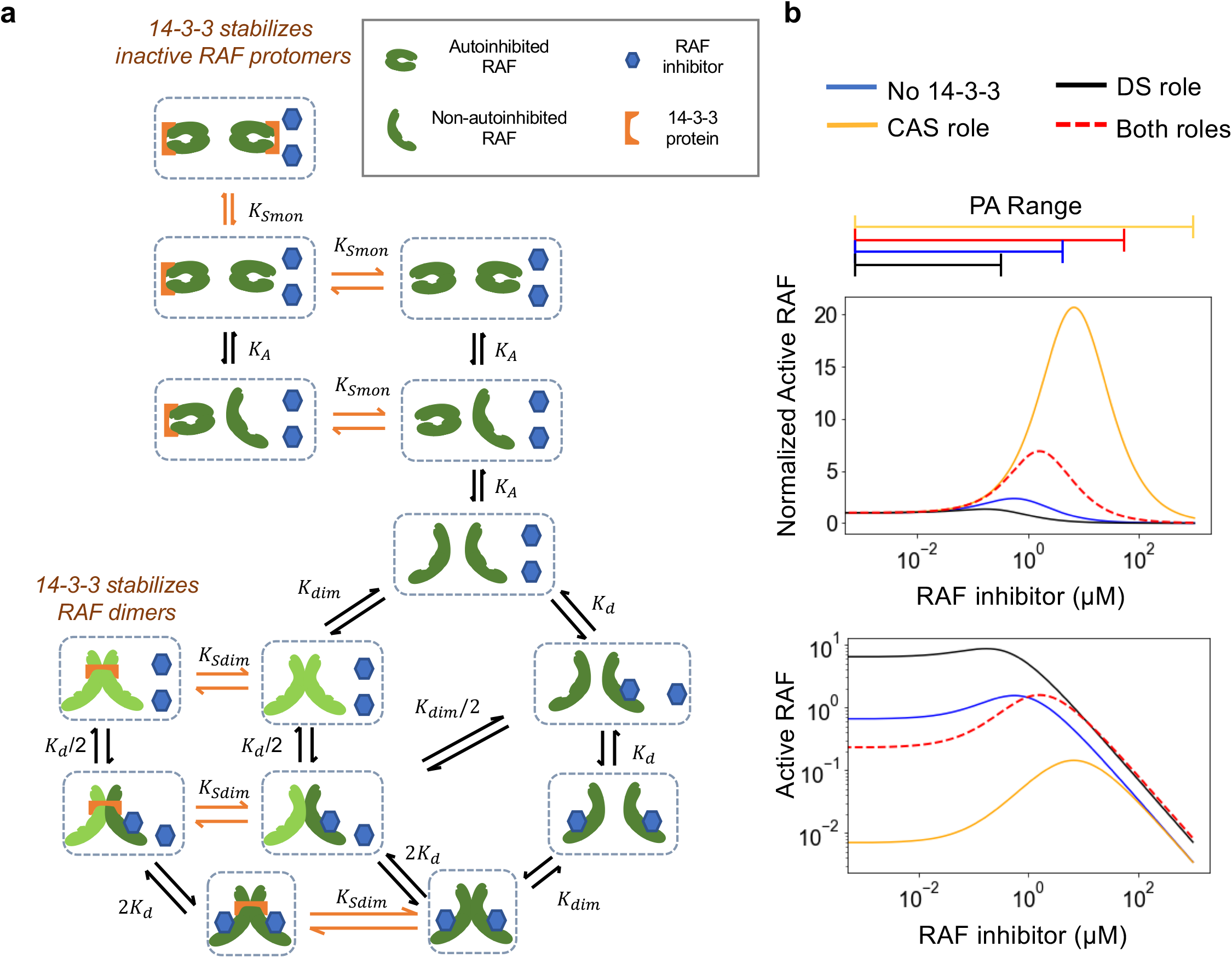
14-3-3 proteins promote paradoxical activation by stabilizing the autoinhibited form of RAF kinases. **a** Schematic of the modeled mechanism of 14-3-3 protein stabilization of the RAF autoinhibited form and the RAF dimer. **b** Predicted dose responses for the RAF autoinhibition model with 14-3-3 stabilizing (or not stabilizing) the autoinhibited form and/or the RAF dimer.

Modeling both roles for 14-3-3 suggests that 14-3-3 proteins should potentiate PA (**Fig. 3b**). To evaluate whether this was due to stabilization of the autoinhibited form, due to stabilization of dimers, or due to both, we considered simplified models where the role of 14-3-3 was limited to only conformational autoinhibition stabilization (CAS), or was limited to only dimer stabilization (DS). By studying these two limiting cases, we found that CAS widened PA to a greater extent than the case where both roles of 14-3-3 were considered. We also found that DS not only failed to potentiate PA, but that DS also reduces PA, thus suggesting that CAS is the dominant mechanism by which 14-3-3 proteins will potentiate PA.

We compared the range of conditions under which PA could occur when 14-3-3 was present and when it was absent. We found that PA is an even more robust outcome of RAF autoinhibitory regulation when 14-3-3 proteins are considered, in that the set of parameters for which PA can result is larger than the set of parameters for which PA can result when no 14-3-3 protein is present (**Supplementary Fig. 2**). Although 14-3-3 proteins promote increased PA, our analysis finds a complex, non-monotonic relationship between the expression level of 14-3-3 with PA magnitude, and between the expression level of 14-3-3 with PA range (**Supplementary Fig. 3**).

### Experiments: 14-3-3 overexpression amplifies paradoxical activation

We experimentally tested the model-based hypothesis that increased expression of 14-3-3 proteins can potentiate PA. Among the seven 14-3-3 encoding genes, we choose *YWHAZ* (which codes for 14-3-3ζ protein) for experimental analysis based on literature spanning several decades and covering biochemical and functional assays connecting this 14-3-3 gene to the RAF activation and inactivation cycle ^20, 47, 48, 49, 50, 51^.

We transfected a *YWHAZ* expression construct into RAS mutant cells. We observed elevated basal signaling, possibly reflecting the promotion of RAF dimers by 14-3-3 (**Fig. 4a**). We then performed a vemurafenib dose response using SK-MEL-2 human melanoma cells, which are BRAF wild-type and harbor an oncogenic *NRAS* Q61R mutation. These cells have previously been utilized to study PA ^18^, and we reproduce PA with vemurafenib here (**Fig. 4b,c**). We observed that SK-MEL-2 cells transfected to express additional, exogenous, 14-3-3ζ displayed significant widening of the PA response compared to the mock transfected cells, consistent with model predictions (**Fig. 4b,c**). In other words, vemurafenib was observed to cause an increase in ERK phosphorylation above the baseline level observed with no vemurafenib. Additionally, the range of vemurafenib doses that showed elevated ERK phosphorylation above baseline was larger in 14-3-3ζ transfected cells than in the mock transfected cells.

**Fig. 4:**
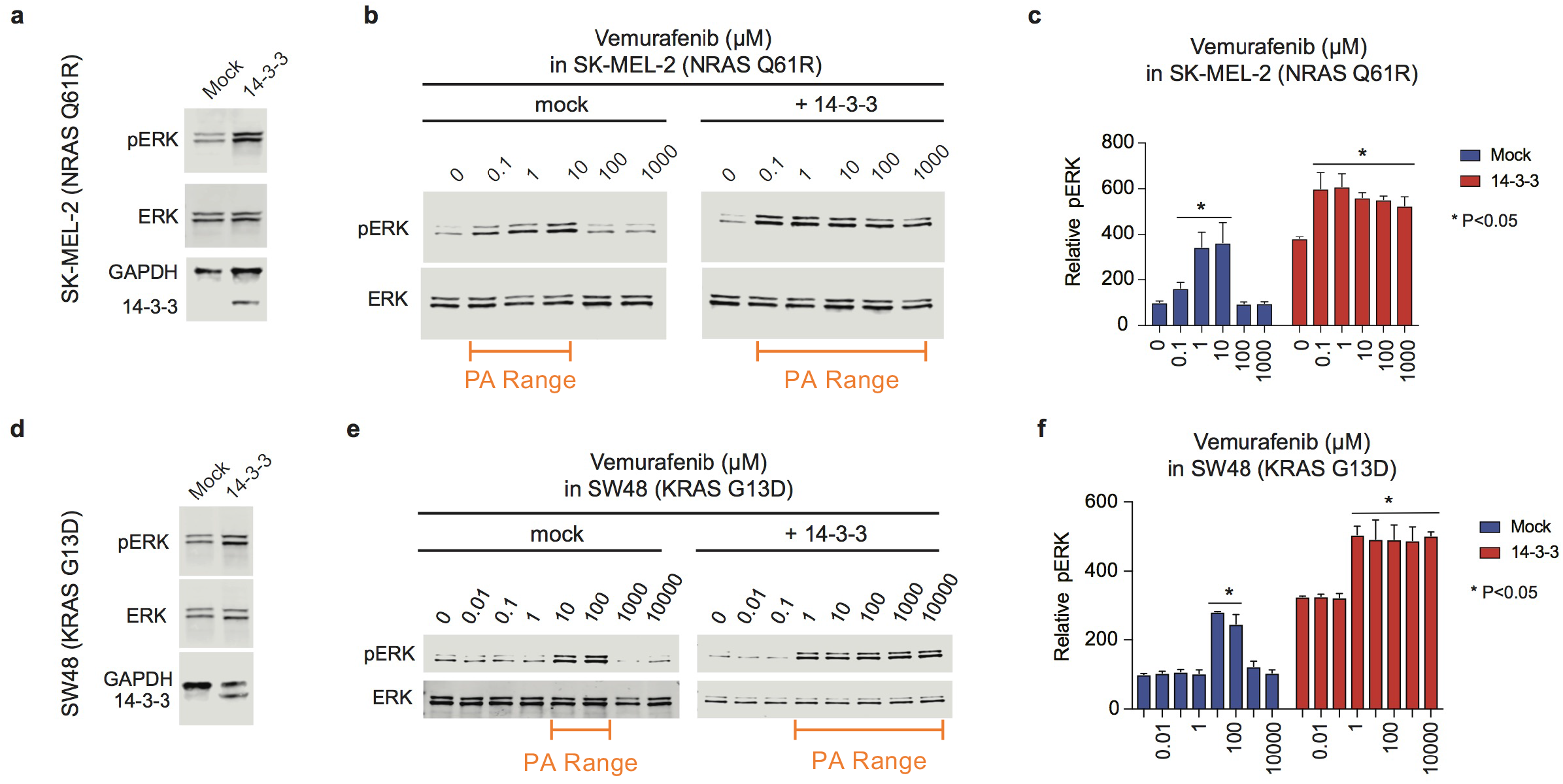
Increased 14-3-3 expression can potentiate paradoxical activation. **a** Immunoblot of SK-MEL-2 cells transfected with 14-3-3 zeta or mock transfected as part of vemurafenib treatment experiments. **b** 14-3-3 and mock transfected SK-MEL-2 cells were treated with increasing doses of vemurafenib, as indicated, for a period of two hours. Cells were then lysed and characterized with immunoblots. Results are representative of three independent experiments. **c** Densitometry-based quantification of the ratio of phosphorylated ERK to total ERK for the indicated doses of vemurafenib from three independent assays. The quantified data are means ± SD. **d** Immunoblot of SW48 KRAS-G13D cells transfected with 14-3-3 zeta or mock transfected as part of vemurafenib treatment experiments. **e** 14-3-3 and mock transfected SW48 KRAS G13D cells were treated with increasing doses of vemurafenib, as indicated, for a period of two hours. Cells were then lysed and characterized with immunoblots. Results are representative of three independent experiments. **f** Densitometry-based quantification of the ratio of phosphorylated ERK to total ERK for the indicated doses of vemurafenib from three independent assays. The quantified data are means ± SD. For panels (c) and (f), one-way ANOVA followed by post-hoc Tukey’s test for multiple comparisons was performed to evaluate differences between drug treated with vehicle treated. * indicates a significant difference with a P value of less than 0.05.

To evaluate the generality of this result, we also performed the same experiments in SW48 colorectal cancer cells that had been engineered to express the *KRAS* G13D mutation. Relative to SK-MEL-2, this cell line introduces a different cancer type, a different *RAS* gene being mutated, and a different hotspot *RAS* mutation. After transfection with 14-3-3ζ (**Fig. 4d**) we treated with vemurafenib. We again observed that increased 14-3-3 expression resulted in PA that spans a wider range of drug concentrations (**Fig. 4e,f**), again consistent with the behavior predicted by our mathematical model.

### Experiments: 14-3-3 overexpression increases paradoxical activation in third generation RAF inhibitors

Ongoing work in RAF inhibitor drug development aims to develop RAF inhibitors where PA does not limit their utility ^3, 4, 5^. These third generation RAF inhibitors should be less prone to dimerization induced resistance, may cause less side effects, and some may even be useful for cancers with a *RAS* mutation. One strategy has been to develop drugs that target both monomeric and dimeric RAF kinases with high affinity and with no or minimal negative cooperativity ^4, 5^. These “pan-RAF inhibitors,” such as LY3009120 and TAK-632, are described to have less PA and for the inhibitory response to occur with a lower quantity of drug ^4, 5^. However, our modeling suggests that 14-3-3 overexpression can stabilize autoinhibitory conformational dynamics to potentiate PA. We therefore hypothesized that 14-3-3 overexpression would counteract the favorable PA profile of pan-RAF third generation RAF inhibitors.

We tested this hypothesis in SK-MEL-2 cells. We transfected with 14-3-3ζ or mock transfected these cells (**Fig. 5a**). We then treated with LY3009120. We detected modest PA for mock transfected SK-MEL-2 cells treated with LY3009120 (**Fig. 5b,c**). This is consistent with studies of these same inhibitors in this same cell line in a previous work ^18^. In SK-MEL-2 cells that express additional 14-3-3ζ, we detected much more robust PA (**Fig. 5b,c**). In our similar evaluation of TAK-632 we found that 14-3-3ζ transfected cells (**Fig. 5d**) displayed more robust paradoxical activation compared to mock transfected cells (**Fig. 5e,f**). For both inhibitors, 14-3-3ζ overexpression resulted in a widening of the range of drug concentrations that show PA. Thus, we find that the favorable PA profile of the “pan-RAF” third generation RAF inhibitors is less favorable when 14-3-3 is expressed at a higher level.

**Fig. 5:**
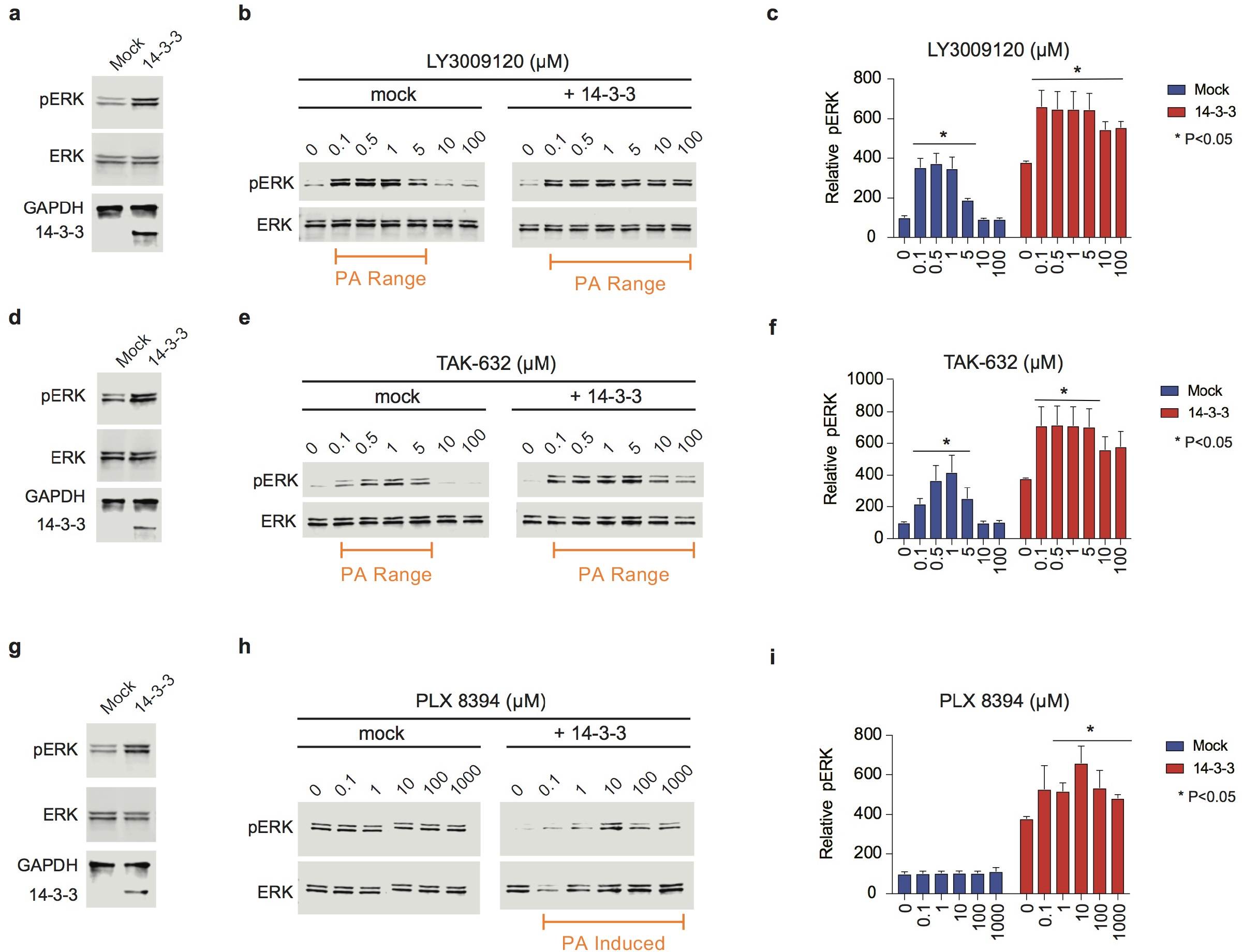
Increased 14-3-3 expression can potentiate paradoxical activation in third generation RAF inhibitors. **a** Immunoblot of SK-MEL-2 cells transfected with 14-3-3 zeta or mock transfected as part of LY3009120 treatment experiments. **b** 14-3-3 and mock transfected SK-MEL-2 cells were treated with increasing doses of LY3009120, as indicated, for a period of two hours. Cells were then lysed and characterized with immunoblots. Results are representative of three independent experiments. **c** Densitometry-based quantification of the ratio of phosphorylated ERK to total ERK for the indicated doses of LY3009120 from three independent assays. The quantified data are means ± SD. **d** Immunoblot of SK-MEL-2 cells transfected with 14-3-3 zeta or mock transfected as part of TAK-632 treatment experiments. **e** 14-3-3 and mock transfected SK-MEL-2 cells were treated with increasing doses of TAK-632, as indicated, for a period of two hours. Cells were then lysed and characterized with immunoblots. Results are representative of three independent experiments. **f** Densitometrybased quantification of the ratio of phosphorylated ERK to total ERK for the indicated doses of TAK-632 from three independent assays. The quantified data are means ± SD. **g** Immunoblot of SK-MEL-2 cells transfected with 14-3-3 zeta or mock transfected as part of PLX8394 treatment experiments. **h** 14-3-3 and mock transfected SK-MEL-2 cells were treated with increasing doses of PLX8394, as indicated, for a period of two hours. Cells were then lysed and characterized with immunoblots. Results are representative of three independent experiments. **i** Densitometrybased quantification of the ratio of phosphorylated ERK to total ERK for the indicated doses of PLX8394 from three independent assays. The quantified data are means ± SD. For panels (c), (f), and (i), one-way ANOVA followed by post-hoc Tukey’s test for multiple comparisons was performed to evaluate differences between drug treated with vehicle treated. * indicates a significant difference with a P value of less than 0.05.

### Experiments: 14-3-3 overexpression creates paradoxical activation in the “Paradox Breaker” third generation RAF inhibitor

Another class of third generation inhibitors, the “paradox breakers,” includes PLX8394. The strategy used to develop the paradox breakers involved finding compounds with a large separation between the dose of inhibitor that inhibits 50% of the pERK signal within a BRAF V600E mutant cell and the dose that promotes 50% of the maximal increase in pERK within a RAS mutant cell ^3^. Based on our analysis of autoinhibitory-driven PA and the role of 14-3-3 proteins, we hypothesized that increased 14-3-3 expression could result in large magnitude PA for this paradox breaker in conditions where it otherwise would not display PA.

To test our hypothesis, we transfected SK-MEL-2 cells to ectopically express 14-3-3ζ (**Fig. 5g**). We then treated these cells with a wide range of doses of PLX8394 (**Fig. 5h,i**). Consistent with previously described observations ^3, 18, 52^, we did not detect PA when we looked at the dose response of PLX8394 in the mock transfected, parental SK-MEL-2. However, with elevated 14-3-3 expression we observed significant PA, consistent with our model-based hypothesis.

### Experiments: 14-3-3 overexpression amplifies RAF-inhibitor induced dimerization

We investigate the effects of 14-3-3 on RAF-inhibitor induced RAF dimerization by bimolecular luciferase complementation. We utilized a previously developed bimolecular luciferase complementation assay that has been combined with integrated Cre-recombinase mediated cassette exchange; this implementation has been named ReBiL for recombinase-enhanced BiLuciferase complementation ^53^ (**Fig. 6a**). Utilizing *RAS* wild-type U2OS cells that have previously been engineered for ReBiL and harbor a floxed, chromosomal, accepter site for efficient generation of stable cell clones that carry both necessary split luciferase fusions, we created a clone with a stable n-terminal split luciferase appended to BRAF and a c-terminal split luciferase appended to CRAF. We then transfected these cells with 14-3-3ζ and treated with RAF inhibitors. We observed a small, statistically significant, increase in cellular proliferation within vemurafenib, LY3009120, and TAK-632 treated cells that had also been transfected with 14-3-3ζ, but not in mock transfected cells treated with the same drugs (**Fig. 6b**). We also quantified cell viability under these same conditions using CellTiter-Glo. Within the time frame of our assays, we observed a RAF-inhibitor induced increase in proliferation that was on top of the baseline increase in proliferation for 14-3-3 transfected cells (**Supplementary Fig. 4**).

**Fig. 6:**
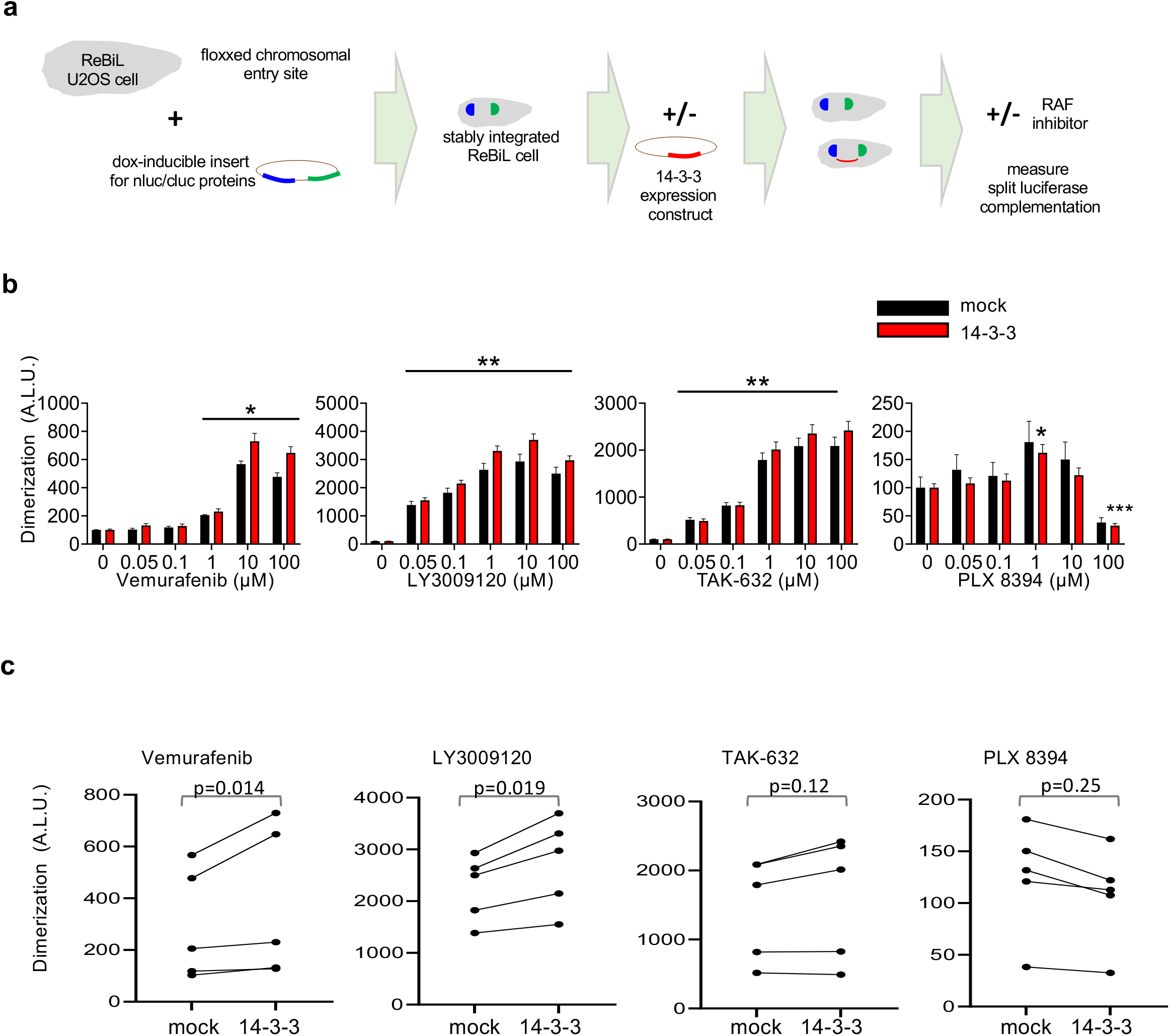
Analysis of RAF inhibitor induced dimer by ReBiL split luciferase complementation. **a** Schematic of the ReBiL assay and its application in these experiments. **b** Luminescent signals from nluc-BRAF, cluc-CRAF U2OS cells treated with the indicated doses of the indicated RAF inhibitors. The marked stars show significant enhancement relative to DMSO baseline. **c** Comparison between luminescence values between mock and 14-3-3 transfected conditions for the same data in (b), but now not normalized to within condition DMSO. Paired t-tests over all doses of the drug show that 14-3-3 transfection led to a statistically significant increase in RAF dimers for Vemurafenib (p=0.01) and LY3009120 (p=0.02) but not for TAK-632 (p=0.06) or PLX8394 (p=0.3). Data shown are from a representative experiment out of two independent biological duplicates. All the panels show mean and ±SD for four replicates in each condition. P-value scheme: * represents a value less than 0.05, ** represents <0.01, *** represents <0.001.

We then investigated RAF-inhibitor induced dimerization. Both 14-3-3ζ and mock transfected cells demonstrated statistically significant increases in dimerization (as suggested by increased luciferase split complementation) relative to the respective no drug condition upon treatment with vemurafenib, LY3009120, and TAK-632 (**Fig. 6c**). For PLX8394, a statistically significant increase in dimerization relative to the respective no drug condition was only observed at a single point of 14-3-3ζ (but not mock) transfected cells. We then evaluated the apparent additional increase in dimerization for 14-3-3ζ transfected cells relative with paired T-tests. We observed statistically significant increases in dimerization within the 14-3-3ζ cells (relative to the mock cells) for vemurafenib and LY3009120 (**Fig. 6d**).

### Experiments: Increased 14-3-3 expression promotes resistance of RAS mutant cells to RAF inhibitors

With the ability to drive increased levels of RAF signaling despite a RAF inhibitor being present, we hypothesize amplified expression of 14-3-3 proteins will emerge as a mechanism that promotes the outgrowth of secondary, RAS-mutant, tumors ^40^ when third generation RAF inhibitors make it to clinical use. To investigate, we propagated SK-MEL-2 cells in media supplemented with PLX8394, media supplemented with vemurafenib, and regular growth media for a period of 21 days (**Fig. 7a**). We measured gene expression in cell lysates collected over the course of 21 days by RNA-seq. We focused on expression levels of the 14-3-3 family of genes. Notably, a statistically significant trend toward increased expression was observed for *YWHAZ* in both the PLX8394 and vemurafenib treated cells (**Fig. 7b,c**). No consistent trend was observed for the other genes that encode 14-3-3 family proteins (**Fig. 7b**). We performed quantitative RT-qPCR to measure *YWHAZ* transcript levels within the same lysates (**Fig. 7d**). By day seven and remaining through the experiment, these measurements also detected a reproducible and statistically significant increase in *YWHAZ* transcript for both vemurafenib and PLX8394 treated cells, thus confirming the observation from the RNA-seq data.

**Fig. 7:**
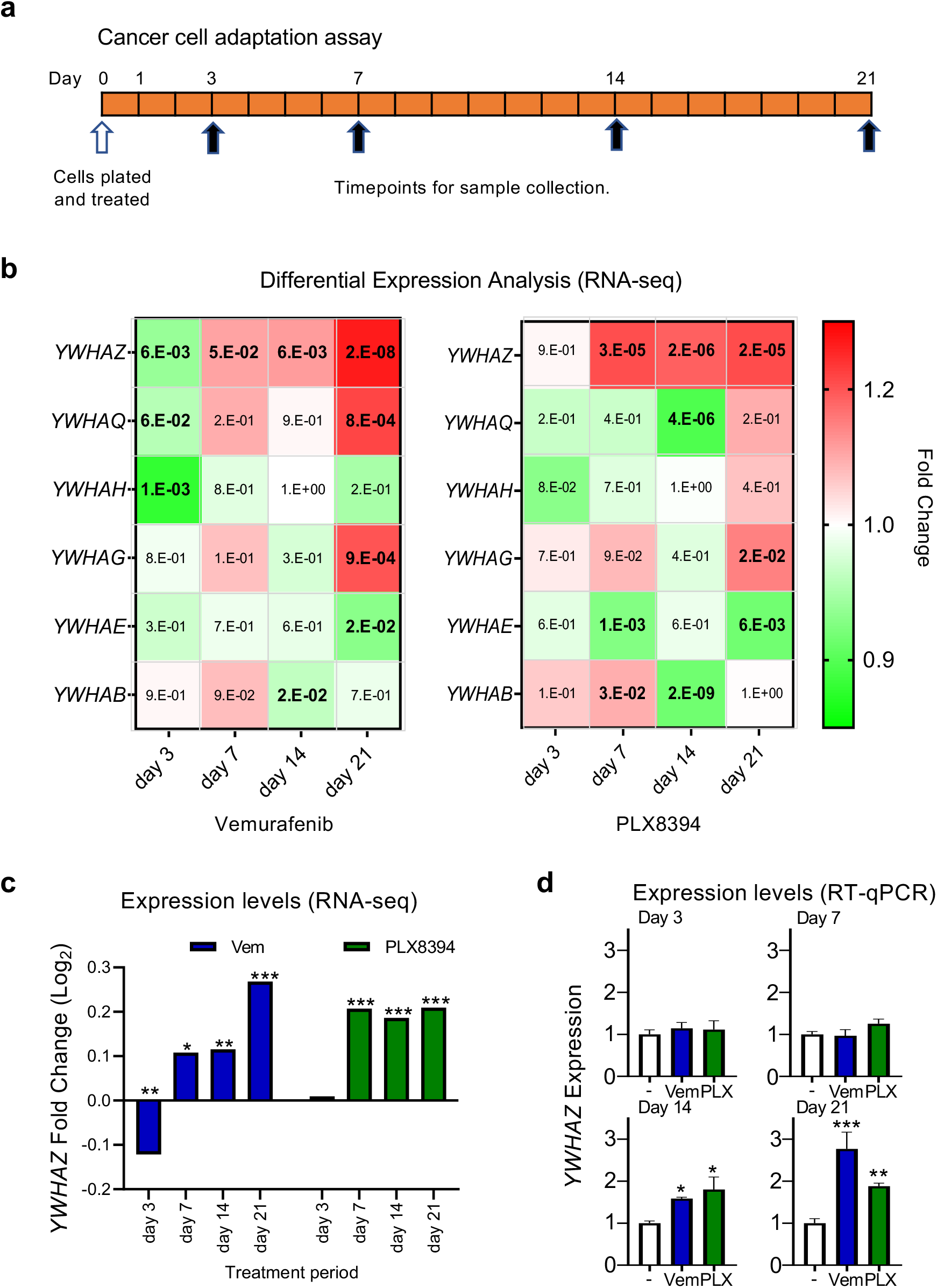
Evaluation of 14-3-3 family gene expression after prolonged treatment with a RAF inhibitor. **a** NRAS mutant SK-MEL-2 cells were treated for 21 days with vemurafenib, PLX8394, or DMSO. At the specified timepoints, samples were saved for gene expression measurements. **b** RNA-seq was performed on the samples indicated in a to evaluate changes in the expression of 14-3-3 family genes. The inset numbers represent adjusted p-value for differential expression of each gene in treated samples relative to respective DMSO control samples. **c** Fold change in *YWHAZ* expression after treatment with vemurafenib and with PLX8394. **d** Relative expression levels measured by RT-qPCR of *YWHAZ* for vemurafenib and PLX8394 treated cells relative to DMSO treated cells at the same time point, as measured by RT-qPCR. p-values shown in (d) were calculated using comparative Ct method. P-values: * represents a value less than 0.05, ** represents <0.01, *** represents <0.001.

### Reconciliation of modeling and experimentation

The experimental work, utilizing three different cell lines, show an activation of the baseline signaling upon increased 14-3-3 expression (Fig. 4a,d, Fig. 5a,d,g, and Supplementary Fig. 4). Comparisons between this observation in baseline conditions with the results of *in silico* model of the RAF activation mechanism (Fig. 3A) for similar baseline conditions (Supplementary Fig. 3c) lead us to conclude that the RAF and 14-3-3 system operates in a part of parameter space where the effects of dimer stabilization are stronger than those of conformational autoinhibition stabilization at baseline. Additionally, the model-informed analysis suggests the baseline level of 14-3-3 that is free to interact with RAF is limiting (Supplementary Figure 3c-f). The key qualifier in the preceding sentence is “free”, as a 14-3-3 protein interacting with another protein would not at that instant also be free to interact with RAF With the many binding partners of 14-3-3 proteins, the quantity of free 14-3-3 is likely to be much smaller than the absolute quantity of 14-3-3 within the cell.

These three cell lines also displayed enhanced PA upon increased 14-3-3 expression (Figs. 4,5,S4). Above, we conclude that dimer stabilization has a stronger effect than conformational autoinhibition stabilization at baseline. However, dimer stabilization acts to counteract PA and conformational autoinhibition stabilization works to amplify PA (Fig. 3b). Although this may superficially appear to be a conflict, mathematical modeling of these mechanisms reveals that it is fully consistent with the mechanism for a large part of parameter space (Supplementary Fig. 3e,f). Thus, we conclude that conformational autoinhibition stabilization by 14-3-3 proteins plays a critical role in the potentiation of PA.

## Discussion

Our mathematical analysis suggests that the conformational regulation of RAF kinase activity combined with conformational dependent dimerization is a critical mechanism that drives PA. Although conformational autoinhibition has been recognized as an important component of the entire PA phenomenon ^15, 27^ it appears to have been underappreciated as a mechanism capable of driving PA. Supporting this assertion is that conformational autoinhibition has neither been discussed as a motivation for the development of third generation RAF inhibitors ^3, 4, 5^ nor has it been included in recent mathematical analyses of PA and of RAF signaling ^38, 39, 54^. This is notable, as both drug development and mathematical analyses pay considerable attention to conformations within the kinase domain (i.e. whether the alpha-C helix and DFG motif are in the “in” or “out” conformations) ^3, 4, 5, 38, 39^. That suggests the concept of structural and conformational factors on PA is not foreign to drug developers and theoretical biologists, but that they may have limited their attention to the small changes in the kinase domain that have dominated the recent literature at the expense of also considering the large, autoinhibitory, conformational changes.

Although we believe that conformational autoinhibition is a critical step in PA that can help explain PA and alleviate ongoing confusion in the field we do not believe it is the only process that contributes to PA. Other processes, like negative allostery (or negative cooperativity) for inhibitor binding ^18^, preferential binding to the different RAF proteins and/or mutant RAF ^2^, altered dimerization affinity for drug-bound RAF proteins ^30, 38, 39^, phosphorylation changes ^23^, scaffold proteins ^24^, allosteric trans-activation ^21^, and RAS nanoclusters ^26^ may all further tune the response to inhibitor, including in drug-specific manners. The possible contributions of negative cooperativity and of drug-induced increases in dimerization affinity have previously received a theoretical treatment ^38^. We believe that ours is the first theoretical analysis of autoinhibitory conformation and its impact on PA.

With respect to 14-3-3 proteins, our mathematical analysis revealed that 14-3-3 protein overexpression could potentiate PA. This includes for agents that normally display minimal to no PA, like the third generation RAF inhibitors. Our experiments robustly detect PA to occur for a larger range of RAF inhibitor concentrations, consistent with our model. Our use of downstream ERK phosphorylation as a readout of RAF kinase activity may introduce a saturable readout of RAF kinase activity, potentially explaining why we can robustly see increases in PA range (which only requires a monotonic relationship between RAF kinase activity and ERK phosphorylation) but not PA fold change (for which the downstream observable would need to be linear as a function of RAF kinase activity for it to be directly measurable). The ability to detect an increase in the range of drug concentrations that display PA would not be limited by saturation and would only require a monotonic relationship between RAF kinase activity and ERK phosphorylation, which seems reasonable to assume. 14-3-3 proteins are promiscuous and pleiotropic, so it is possible that part of the effects observed with 14-3-3 transfection follow from other activities of 14-3-3 proteins; it is not possible to rule-out unknown alternative mechanisms involving 14-3-3 proteins. At minimum, our mathematical analysis yielded a novel hypothesis that led to the empirical observation that 14-3-3 protein amplification potentiates PA. With the ability to drive increased levels of RAF signaling despite a RAF inhibitor being present, we hypothesize amplified expression of 14-3-3 proteins will emerge as a mechanism of resistance and toxicity should third generation RAF inhibitors make it to clinical use.

## Materials and Methods

### Mathematical models and analysis

We focus on steady-state levels of the different states in which RAF can exist, as portrayed in the diagrams for each model. Between any two states an equilibrium relationship can be expressed as the ratio of abundances in the two states. Conservation of total protein quantities and zero value of total Gibbs free energy change at equilibrium both provide mechanisms to algebraically combine these expressions. We thereby derive algebraic expressions that relate the relative abundance of the RAF within its different monomeric and dimeric states. We perform algebraic manipulations and derive analytic solutions which we cross-check using Mathematica software (Wolfram Research). We perform numerical evaluations of these relationships, and generate plots of these equations using Python packages including numpy, scipy and matplotlib.

### Cell culture and transfection

SK-MEL-2 cells were purchased from American Type Culture Collection (ATCC). SW48 cells with the G13D genotype were obtained from Horizon Discovery. The DNA expression plasmid used for 14-3-3, 1481 pcDNA3 flag HA 14-3-3ζ, was a gift from William Sellers (Addgene plasmid # 9002; http://n2t.net/addgene:9002; RRID:Addgene_9002). Cells were grown in EMEM (SK-MEL-2) or RPMI (SW48) containing 10% fetal bovine serum (FBS), penicillin (100 U/ml), streptomycin (100 μg/ml), and l-glutamine (2 mM). Cells were cultured in 10cm adherent culture dishes (VWR) and incubated at 37°C in 5% CO_2_. At time point zero, cell media was changed to the cell line’s respective media containing 10% fetal bovine serum (FBS) devoid of antibiotics. 24 hours later cells were transfected with empty packaged lipofectamine, or 5ug of 14-3-3ζ expression plasmid DNA utilizing lipofectamine 2000 (Thermo Fisher Scientific) following manufacturers protocol. Cells were incubated for 24 hours and then were treated with RAF inhibitors at increasing doses for 2 hours. Cells were then prepared for Western blot analysis. All drugs were suspended and stored in DMSO, and all drug treatment groups carried the same amount of vehicle (DMSO).

### Western blotting

Cell lysates were generated using radioimmunoprecipitation assay buffer [150 mM NaCl, 1% NP-40, 0.5% sodium deoxycholate, 0.1% SDS, 50 mM tris (pH 8.0)] containing protease and phosphatase inhibitor cocktail (Cell Signaling Technology) and incubated on ice for 1 hour, vortexing every five minutes. The total protein concentration was determined by Pierce Protein assay (Thermo Fisher Scientific). Protein samples (5μg) were resolved by electrophoresis on 12% SDS–polyacrylamide gels and electrophoretically transferred to polyvinylidene difluoride (PVDF) membranes (Millipore Corporation) for 20 min at 25 V. The blots were probed with the mouse anti-phospho-ERK antibody (675502, Biolegend) and rat anti-total-ERK antibody (686902) overnight at 4 degrees Celsius. Blots were washed and probed with goat-anti-mouse Dylight 800 secondary antibody and goat-anti-rabbit AlexaFlour 680 antibody for 1 hour at room temperature. The protein bands were visualized using the Licor CLx Odyssey imaging station (Licor Biosystems). Comparative changes were measured with Licor Image Studio software.

### RAF inhibitor resistance assay

SK-MEL-2 cells were grown in EMEM (HyClone) supplemented with 10% fetal bovine serum (Atlanta Biologicals) and Penicillin-Streptomycin (Gibco). The cells were treated with the vehicle (DMSO), PLX8394 (10μM), or Vemurafenib (10 μM) for 0, 3, 7, 14, or 21 days. Total RNA from SK-MEL-2 human melanoma cells was harvested for RT-qPCR and RNA sequencing using the E.Z.N.A Total RNA Kit I (Omega Bio-Tek). The quality and quantity of the total RNA was assessed using NanoVuePlus Spectrophotometer (GE Healthcare) in sample preparation, and more precisely evaluated prior to sequencing using Agilent TapeStation 4200. RNA-Seq libraries were prepared with 500 ng total RNA using the TruSeq stranded mRNA Sample Preparation Kit according to the manufacturer’s protocol (Illumina). RNA-seq libraries were multiplexed, normalized and pooled for sequencing. The libraries were sequenced on the HiSeq 4000 system (Illumina) at single read 50bp. Image analysis and base calling were done with Illumina CASAVA-1.8.2. on HiSeq 4000 system and sequenced reads were quality-tested using FASTQC. The GEO accession number for the RNAseq data is GSE179932. Reverse transcription was performed with the iScript Supermix cDNA Synthesis Kit (Bio-Rad). Quantitative polymerase chain reaction was performed by using the PerfeCTa SYBR Green FastMix (Quanta BioSciences) and CFX96 Touch Real-Time PCR Detection System (Bio-Rad). Transcripts were amplified using gene-specific primers for YWHAZ (F: 5’-CAACAAGCATACCAAGAAG-3’; R: 5’-TCATAATAGAACACAGAGAAGT-3’) and GAPDH (F: 5’-ACCACAGTCCATGCCATCAC-3’; R: 5’-GCTTCACCACCTTCTTGATG-3’). The comparative Ct method was used for relative quantification for gene expression data analysis in the quantitative RT-qPCR analysis.

### RNA-Seq analysis

Sequenced reads were quality-tested using FASTQC (v0.11.8) ^55^ and mapped to the hg19 human genome using the STAR aligner (v2.5.3a) ^56^ with default parameters. Raw or TPM (Transcripts Per Kilobase Million) gene expression was quantified across all the exons of RefSeq genes with analyzeRepeats.pl in HOMER (v4.11.1) ^57^, which used the top-expressed isoform as proxy for gene expression. Differential gene expression was performed with the raw gene counts using the R package, DESeq2 (v1.24.0) ^58^, using replicates to compute within-group dispersion. Differentially expressed genes were defined as having a false discovery rate (FDR) <0.05 when comparing two experimental conditions.

### ReBil assay to measure RAF dimerization

The full-length human BRAF and CRAF were cloned to pLi635 ReBiL targeting plasmid ^59^ by Gibson assembly (NEBuilder HiFi E2621L) resulting in pLi814 (nl-BRAF cl-CRAF), which was integrated to U2OS 134-8 HyTK8 via Cre-recombinase mediated cassette exchange (RMCE) ^53^ resulting in U2OS 134-814 ReBiL cell line. U2OS 134-814 ReBiL cells cultured in T25 flask at 37°C 7% CO_2_ incubator with 6 ml DMEM (Corning; 10-013-CV) with 10% (vol/vol) FBS, and 10 μ g/mL ciprofloxacin (Corning; 61-277-RG) were transfected with 7.5 μg pcDNA3 or pcDNA3_flag_HA_14-3-3ζ by Lipofectamine 3000 (Thermo Fisher Scientific) and incubated overnight. Approximately 5,000 transfected ReBiL cells were seeded into two 384-well plates and treated with 100 ng/ml doxycycline to induce nluc-BRAF and cluc-CRAF expression for 24 hours. RAF inhibitors were added at indicated concentrations and incubated for 2 hours. One-Glo (Promega E6120) was added to one plate and CellTiter-Glo 2.0 (Promega G9242) was added to another plate. Luminescent signals were measured by Tecan M200 microplate reader (integration time 0.5 sec at 26 °C). Significance analysis was performed assuming normally distributed variation with a two sided t-test.

## Acknowledgments

We thank Reuben Shaw, Bjorn Lillemeier, Dmitry Lyumkis, Joe Noel, Andrey Shaw, Tony Hunter, Michael Trogdon and members of the Stites lab for helpful conversations and feedback. We thank Nasun Hah and the members of the Salk Next Generation Sequencing Core for their assistance, guidance, and access to equipment. We thank Amy Cao for help with illustrative graphics.

## Author contributions

G.M and E.S. designed the study. G.M. performed the mathematical and computational analyses. K.A., T.M., L.L., G.M., J.H., G.W. and E.S. designed the experiments. K.A., L.L., and T.M. performed the experiments. G.M., J.Y., M.S., analyzed the genomic data. G.M., M.S., and E.S. determined the needed statistical analyses and G.M., T.M., J.Y. and K.A. performed the statistical analysis. E.S. and G.M. wrote the manuscript with input from the other authors.

---

**Supplementary Fig. 1:**
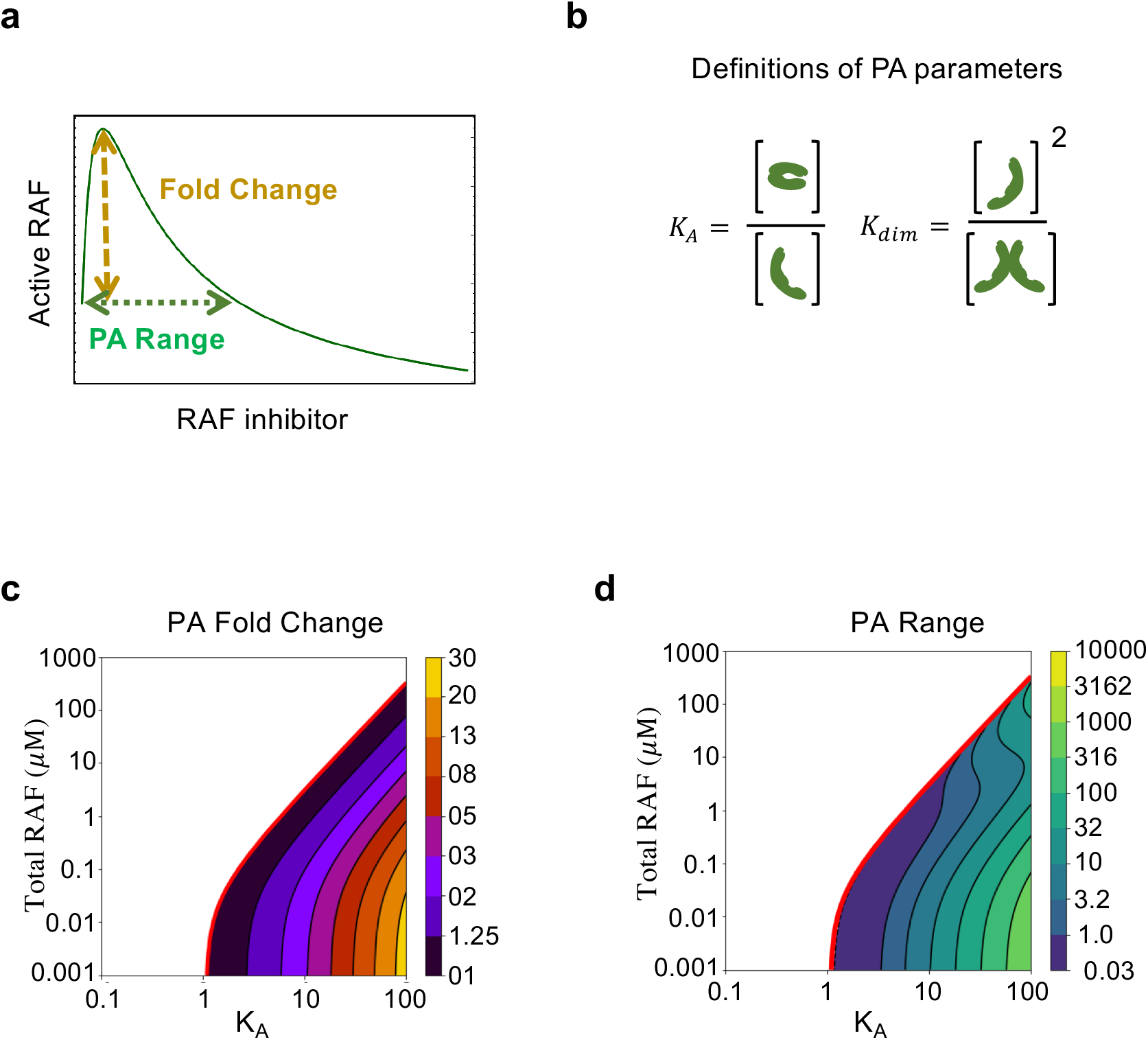
Magnitudes of paradoxical activation as a function of autoinhibition propensity and RAF abundance. **a** Schematic to define two measures of PA: PA Peak Fold Change, which is the ratio of peak intensity to baseline (without drug) intensity of active RAF protomers; and, PA Range, which is the drug concentration at which the inhibitory phase begins. **b** Schematic to define the key equilibrium constants considered in the base model. K_A_, or the autoinhibition equilibrium constant and K_dim_, or the dimerization constant for RAF. **c** Predicted PA Peak Fold Change presented as a function of two key parameters of the autoinhibition model (K_A_ and RAF_rel_). **d** Predicted PA Range, presented as drug concentration relative to drug affinity as a function of a for two key parameters of the model (K_A_ and RAF_rel_).

**Supplementary Fig. 2:**
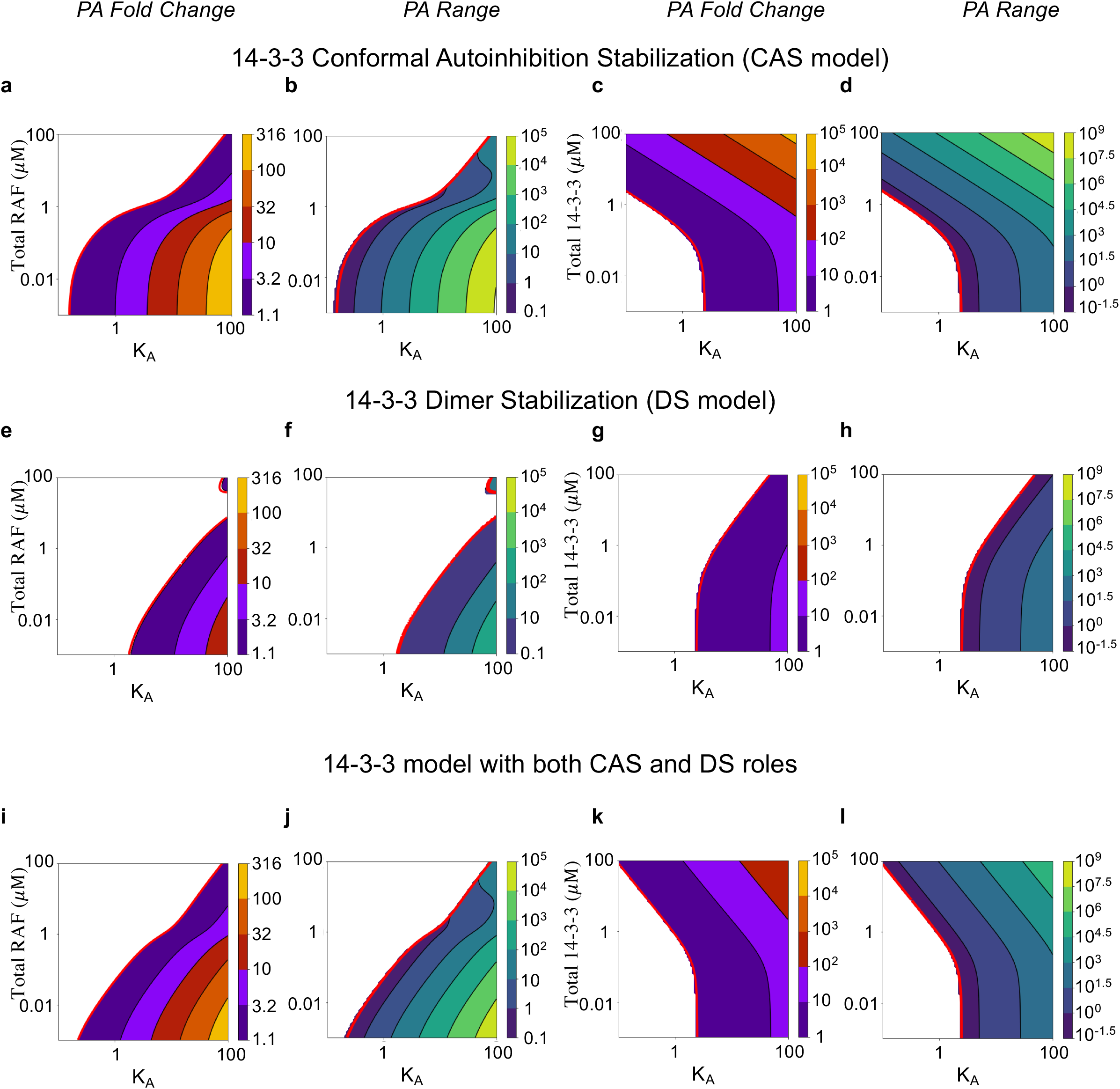
Magnitudes of paradoxical activation as a function of autoinhibition propensity, RAF abundance, and 14-3-3 abundance in a system with 14-3-3 proteins. The panels show contour plots with RAF concentration (a,b,e,f,i,j) or 14-3-3 concentration (c,d,g,h,k,l) on the y-axis, and with the intrinsic autoinhibition parameter K_A_ on the x-axis. The contours represent predicted PA Fold Change (a,c,e,g,i,k) and predicted PA Range (b,d,f,h,j,l). ad Show results in CAS model. e-h Show results in DS model. i-l Show results in 14-3-3 model with both CAS and DS roles.

**Supplementary Fig. 3.**
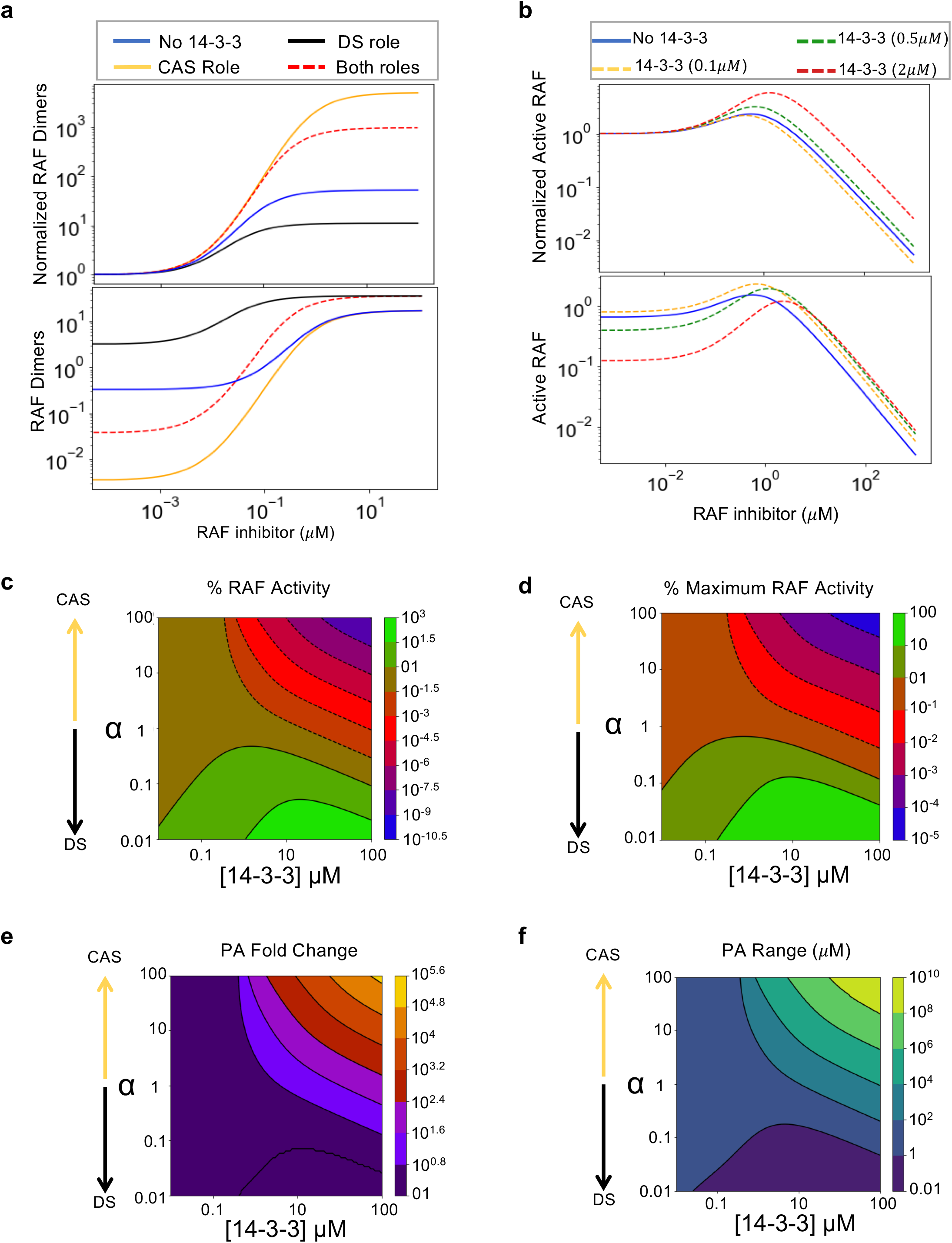
Further predictions with both CAS and DS roles of 14-3-3 interactions. **a** Shows how 14-3-3 proteins would influence total RAF dimers when only the CAS mechanism, only the DS mechanism, or both CAS and DS mechanisms are considered. The RAF dimers are normalized to drug free level (above) and percent of total RAF (below) for the same conditions as Fig. 2b. **b** Shows activation curves as a function of drug for active RAF protomers for increasing levels of 14-3-3 expression. Active RAF is shown normalized to the drug free level (above) and to the percent of total RAF (below) for the same conditions as figures 2b. **c** Presents baseline RAF activity (without drug). **d** Presents percent maximum RAF activity (not normalized to drug-free condition) **e** Presents PA fold change. **f** Presents PA range. For c-f, the x-axis displays for increasing levels of 14-3-3 expression (from 1nM to 100uM). For panels c-f, the y-axis displays varying degrees of the relative strength of DS and CAS mechanisms. This is specified as a meta-parameter *α* which is multiplied to the dissociation of 14-3-3 interaction with RAF dimers (*K_Sdim=α×K_Sdim*) and divided from the dissociation constant of 14-3-3 interaction with inactive RAF monomers (*K_Smon=K_Smon/α*). A small *α* favors strong DS and weak CAS, and a large *α* favors weak DS and strong CAS.

**Supplementary Fig. 4.**
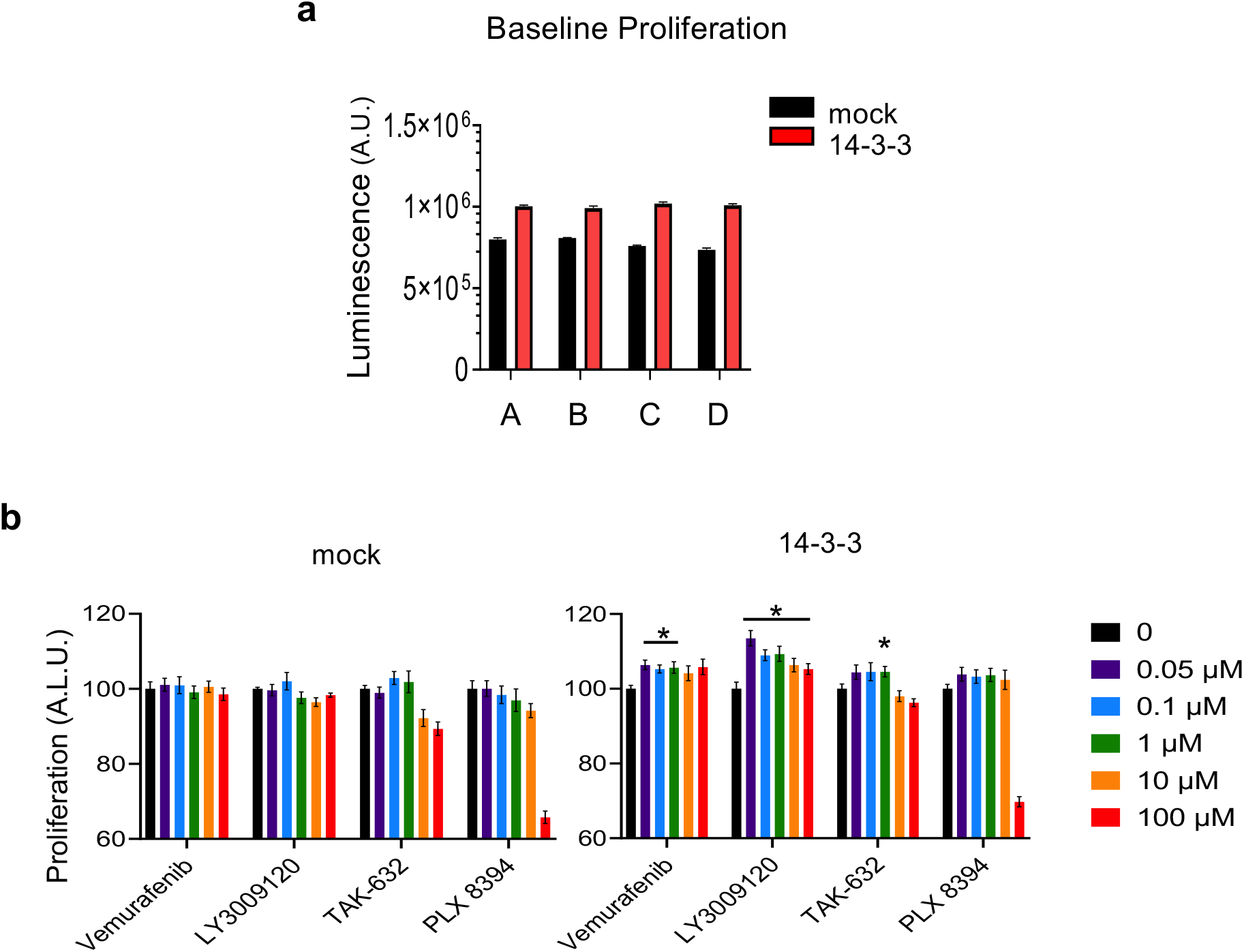
Baseline toxicity in ReBiL cells evaluated via proliferation assay. The nluc-BRAF, cluc-CRAF, stable, U20S cell clones had proliferation measured with CellTiter Glo under mock and 14-3-3 transfected conditions after 2 hours of drug exposure. Samples A, B, C, and D were the stocks used for the study of vemurafenib, LY3009120, TAK-632, and PLX8394, respectively in the ReBiL assay.

